# Trait anxiety impairs reciprocity behavior: A multi-modal and computational modeling study

**DOI:** 10.1101/2024.07.21.604469

**Authors:** Huihua Fang, Rong Wang, Zhihao Wang, Qian Liu, Yuejia Luo, Pengfei Xu, Frank Krueger

## Abstract

Anxiety significantly impacts reciprocal behavior, crucial for positive social interactions. The neurocomputational mechanisms of anxiety’s effects on the core (individual propensity) and peripheral (decision context) factors shaping reciprocity remain unclear. Here, we investigated reciprocity in individuals with low and high trait anxiety using a binary trust game with gain/loss framing, combining computational modeling, eye-tracking, and event-related potentials (ERPs). Our computational model, validated by eye-tracking data, identified four psychological components driving reciprocal behavior: reward, guilt aversion, superiority aversion, and superiority attraction. Regarding the core of reciprocity, trait anxiety diminished both overall reciprocity and specific psychological components like guilt aversion and superiority attraction, irrespective of context. The reduction in guilt aversion was supported by ERP findings showing decreased P2 (selective attention) and increased LPP (emotion regulation) amplitudes in anxious individuals. Regarding the periphery of reciprocity, trait anxiety altered the contextual perception of both superiority aversion and reward. Further, trait anxiety reversed the perception of superiority aversion from gain to loss contexts, a pattern that was linked to the N2 amplitudes (cognitive control). Our findings revealed distinct effects of trait anxiety on core and peripheral factors in reciprocity, offering potential targets for interventions aimed at improving reciprocity in individuals with anxiety disorders.

## 1. Introduction

Reciprocity acts as a crucial glue for interpersonal interactions, playing a pivotal role in cultivating a harmonious and thriving society [1]. As the global economic slowdown intensifies and geopolitical conflicts escalate, anxiety levels among populations have increased significantly [2]. Hence, understanding the impact of anxiety on reciprocity is essential for maintaining social cohesion.

Social representation theory conceptualizes reciprocity decisions as having two main aspects: a core and a periphery [3–6]. The core represents an individual’s inherent reciprocity propensity, which remains stable across different contexts. The periphery reflects an individual’s contextual perception, shaping the decision by taking into account the situational context. Disentangling reciprocity propensity from contextual dynamics continues to be a challenge in decision-making. One effective solution is to systematically adjust the periphery by reframing the decision context while maintaining the same payoff structure (e.g., framing gains vs. losses) [4,7]. This approach enables the assessment of how trait anxiety influences both the core and periphery of reciprocity, clarifying whether these effects stem from the inherent reciprocity propensity or contextual modifications.

In social interactions, reciprocating rather than betraying a partner’s trust helps mitigate negative emotions such as guilt aversion (failure to meet a partner’s reciprocating expectations) and superiority aversion (also known as advantageous inequity aversion: feeling discomfort from receiving more than others). However, this often comes at a cost—sacrificing personal benefits, such as financial rewards, which would be the economically optimal choice [8,9].

While guilt aversion and reward are widely recognized as key components affecting prosocial behaviors, the concept of superiority aversion remains controversial. For example, superiority aversion is not always present; it emerges only when the decision-maker actively chooses an advantageous inequitable distribution but disappears when the distribution is passively received [10]. Furthermore, evidence suggests that people constantly compare themselves with others [11–13], often seeking to increase their own payoff relative to others, demonstrating superiority attraction instead of aversion under certain circumstances [14–16]. Since betrayal typically leads to a more advantageous distribution than reciprocity, it is plausible that individuals exhibit superiority aversion when considering betrayal but show a preference for superiority when evaluating reciprocity. This scenario creates a complex social dilemma for decision-makers, who must balance moral considerations, social comparison, and personal interests.

Recent research has increasingly utilized computational models to explore the complex psychological components driving reciprocity [8,9,17]. Those computational models typically suggest that decision-makers aim to maximize the utility of their choices by balancing economic benefits, such as monetary rewards, against the costs of norm violations, including anticipatory negative emotions like guilt and superiority aversion. However, the extent to which these psychological components reflect the actual psychological processes underlying decision-making has rarely been validated by empirical evidence. Eye-tracking during decision-making, such as in economic games where participants look at payoff structures, can provide direct observation of the decision-maker’s focal points [18–20], allowing to validate whether these focuses align with the estimates derived from the proposed model. For example, decision-makers who are more sensitive to guilt aversion are expected to spend relatively more time looking at those payoff structures that trigger guilt aversion. Combining computational modeling and eye-tracking provides deeper insights into the psychological processes underlying reciprocity, including how factors like anxiety influence these processes.

Research on social anxiety and generalized anxiety disorder has shown diminished generosity [21], cooperation [22], and reciprocity [23–25], suggesting a broad inhibitory effect of anxiety on prosocial behaviors including reciprocity. Individuals with obsessive-compulsive personality disorder (OCPD), which often co-occurs with high anxiety levels, exhibit less guilt aversion in reciprocity decisions during a binary trust game, suggesting that anxiety might attenuate anticipatory feelings of guilt by restricting affective processing and reducing their sense of moral obligation [17]. Additionally, anxious individuals tend to adopt avoidance strategies [26–28] and engage in excessive effortful processing when regulating emotion [29–31]. Overall, this evidence suggests that anxious individuals may mitigate anticipatory aversive feelings like guilt or superiority aversion during reciprocity decisions by actively avoiding negative emotions or reducing attention.

Individuals with high trait anxiety are particularly susceptible to contextual effects [32–34], which can influence the periphery of reciprocity. Previous studies have mainly focused on non-social aspects of decision-making, like risk evaluation in gambling tasks, rather than on tasks involving social interactions that balance social norms and personal economic benefits [32–34]. Although the contextual effects on social reciprocity have been investigated [7], the impact of anxiety on these contextual influences and the underlying psychological components of reciprocity remains unexplored. Anxious individuals rely more on heuristics when making decisions, evidenced by increased brain activity in regions associated with cognitive effort during frame-inconsistent decisions [33,35]. Further, loss compared to gain frames are heuristically perceived as more harmful to others [7,36], and more threatening to one’s own payoff [34]. Therefore, anxiety may alter the perception of other-regarding components, such as guilt and superiority aversion, and self-beneficial rewards in reciprocity, likely involving cognitive control mechanisms.

To understand how anxiety affects reciprocity, it’s essential to investigate the neuropsychological mechanisms by which anxiety influences contextual perceptions and reciprocity propensity. Electroencephalography (EEG), particularly event-related potentials (ERP), can offer significant insights into the temporal dynamics of neural mechanisms. For example, studies have linked P2 with selective attentional allocation [37–40] and N2 with effortful top-down cognitive control in decision-making [41–44]. Although late positive potential (LLP) is often seen as a marker of emotional reactivity [45–49], several studies have linked LPP with emotional regulation and found that an enhanced LPP amplitudes index an increase of cognitive effort in managing emotional responses [50–53]. Specifically, in decisions involving moral conflict, larger LPP amplitudes indicate more cognitive efforts deployed to resolve these conflicts [54–56].

To investigate the neurocomputational mechanisms underlying reciprocity decisions, we formulated four hypotheses. First, based on prior research [8–10,14–17], we hypothesized that the best computational model for reciprocity decisions would consist of four distinct psychological components: reward, guilt aversion, superiority aversion, and superiority attraction. Second, we predicted that individuals more sensitive to these psychological components would spend relatively more time visually attending to the according payoff structure within the binary trust game, reflecting their evaluation processes during reciprocity decisions [18–20]. Third, regarding the core of reciprocity, we expected trait anxiety to attenuate the core of reciprocity across gain and loss frames, given that anxious individuals exhibit reduced reciprocity behavior [23–25]. In particular, we hypothesized that high trait anxiety would reduce guilt and superiority aversion, as anxious individuals tend to adopt avoidance strategies [26–28] and mitigate negative feelings by limiting affective processing and reducing moral obligation [17,57]. We anticipated this process would manifest in the P2 and LPP ERP components, given P2’s association with selective attentional allocation [37–40] and LPP’s connection to emotional regulation [50–53] and moral conflict [54–56]. Fourth, regarding the periphery of reciprocity, we postulated that trait anxiety modulates reciprocity differently under gain and loss contexts, given that anxious individuals are more susceptible to contextual influences [32,34]. We proposed that trait anxiety would differentially modulate other regarding components (i.e., guilt aversion and superiority aversion) and reward sensitivity across varying contextual frames, given that anxious individuals often rely on heuristics in decision-making [33], and that loss frames are generally perceived as more detrimental to others [7,36] and to one’s own economic interests [34]. We predicted these anxiety-related alterations would engage N2 cognitive control mechanisms.

To test our hypotheses, we combined EEG, eye-tracking, and computational modeling in a binary trust game to investigate the neurocomputational mechanisms of reciprocity decisions in individuals with low and high trait anxiety under gain and loss frames (**Fig. 1**). Participants also completed in addition two self-report questionnaires, the Interpersonal Reactivity Index (IRI) measuring aspects of empathy [58] and Machiavellianism (Mach-IV) scale measuring the tendency towards selfishness [59]. Our computational modeling analysis identified four psychological components underlying reciprocity decisions—reward, guilt aversion, superiority aversion, and superiority attraction—which were validated by the eye-tracking analyses. Regarding the core, our findings revealed that trait anxiety reduced reciprocity at the behavioral level and diminished guilt aversion and superiority attraction at the psychological level. At the neural level, trait anxiety attenuated guilt aversion by decreasing P2 amplitude, related to selective attention, and increasing LPP amplitude, associated with effortful emotion regulation. Regarding the periphery, trait anxiety influenced the contextual effects on reward and superiority aversion at the psychological level, although it did not affect reciprocity behaviorally. Specifically, trait anxiety modified the contextual effect on superiority aversion through the N2 cognitive control mechanism at the neural level.

**Figure 1.**
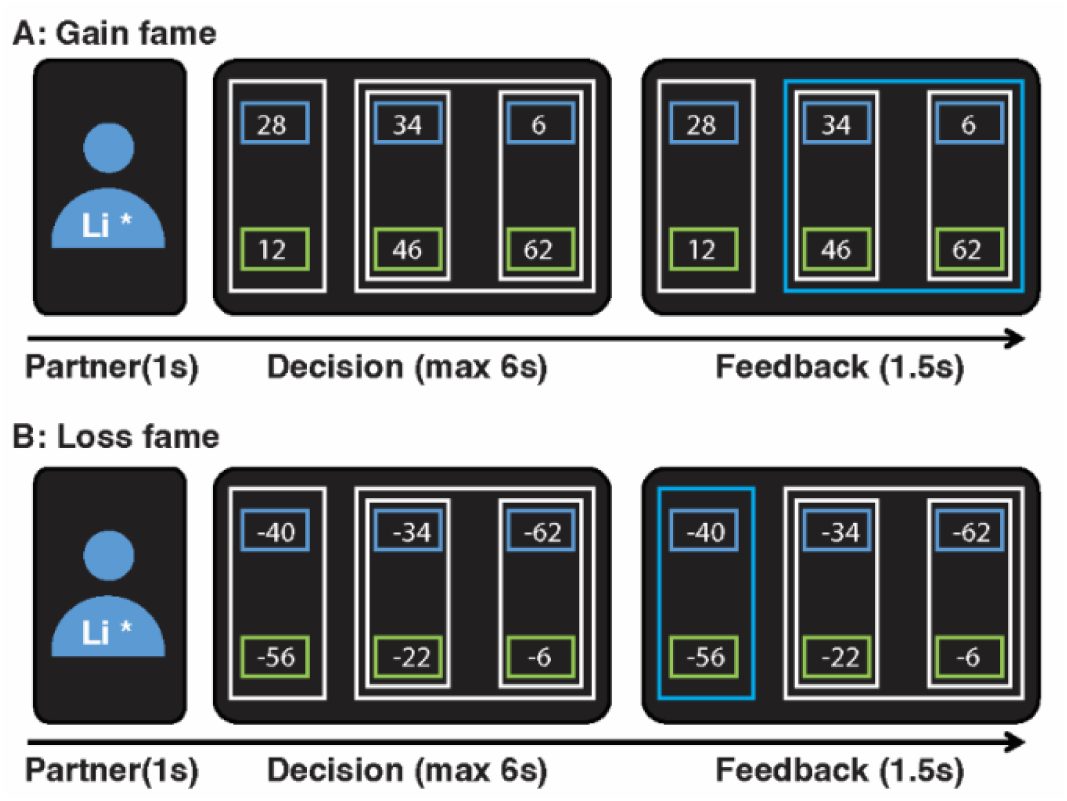
Experimental design. Participants completed a trust game with two distinct frame sessions in a counterbalanced order: **(A)** Gain frame session and **(B)** Loss frame session. The gain frame session started with 0 points while the loss frame session started with 15,000 points. If participants used the same strategies in both frames, their final outcomes would be identical [7]. For each trial session, participants were introduced to a different anonymous partner, represented by an icon with a partially obscured name (i.e., one word in the first name was blocked). Participants then decided whether to “reciprocate” (e.g., in the gain frame, choosing the middle rectangle resulted in a distribution of 46 points for themselves and 34 points for their partner; in the loss frame, it involved a deduction of 22 points for themselves and 34 points for their partner) or “betray” (e.g., in the gain frame, choosing the right rectangle resulted in a distribution of 62 points for themselves and 6 points for their partner; in the loss frame, it involved a deduction of 6 points for themselves and 62 points for their partner) without knowing whether the partner had chosen “trust” (e.g., the larger rectangle containing both the “reciprocate” and “betray” option) or “distrust” (e.g., in the gain frame, the left rectangle resulted in a distribution of 12 points for themselves and 28 points for their partner; in the loss frame, it involved a deduction of 56 points for themselves and 40 points for their partner). Finally, participants received feedback with a blue highlight indicating whether the partner had initially chosen to “trust” (as shown in **A**) or “distrust” (as shown in **B**). Participants were unaware the feedback was randomly generated by the computer. Participants were informed of the rules before the game that if the partner had chosen to “distrust,” the payoff would be distributed accordingly, but if the partner had chosen to “trust,” the payoff would be based on the participant’s decision.

## 2. Results

### 2.1. Computational Modeling of Reciprocity

Computational modeling was employed to unveil the psychological components behind reciprocity as measures with the binary trust game. Seven plausible candidate models (see **Materials and Methods**)— adapted from previous studies [8,9,17] with components including reward, guilt aversion, inequity aversion, superiority aversion, and superiority attraction—were constructed and compared using a stage-wise approach [60–62].

The proposed model 4 (**M4**, pseudo *r*^2^ = 0.359; **Fig. 2A**; **Tab. 1**) outperformed all other candidate models, as indicated by the leave-one-out information criterion (LOOIC) and widely applicable information criterion (WAIC). **M4** included the components of reward sensitivity, guilt aversion, superiority aversion, and superiority attraction, explaining over 93% of the variance in reciprocity rates (**Fig. 2B**). Higher subjective sensitivity to guilt aversion, superiority aversion, and superiority attraction was associated with increased reciprocity rates, while greater sensitivity to reward with lower reciprocity rates (**Fig. 2C**).

**Figure 2.**
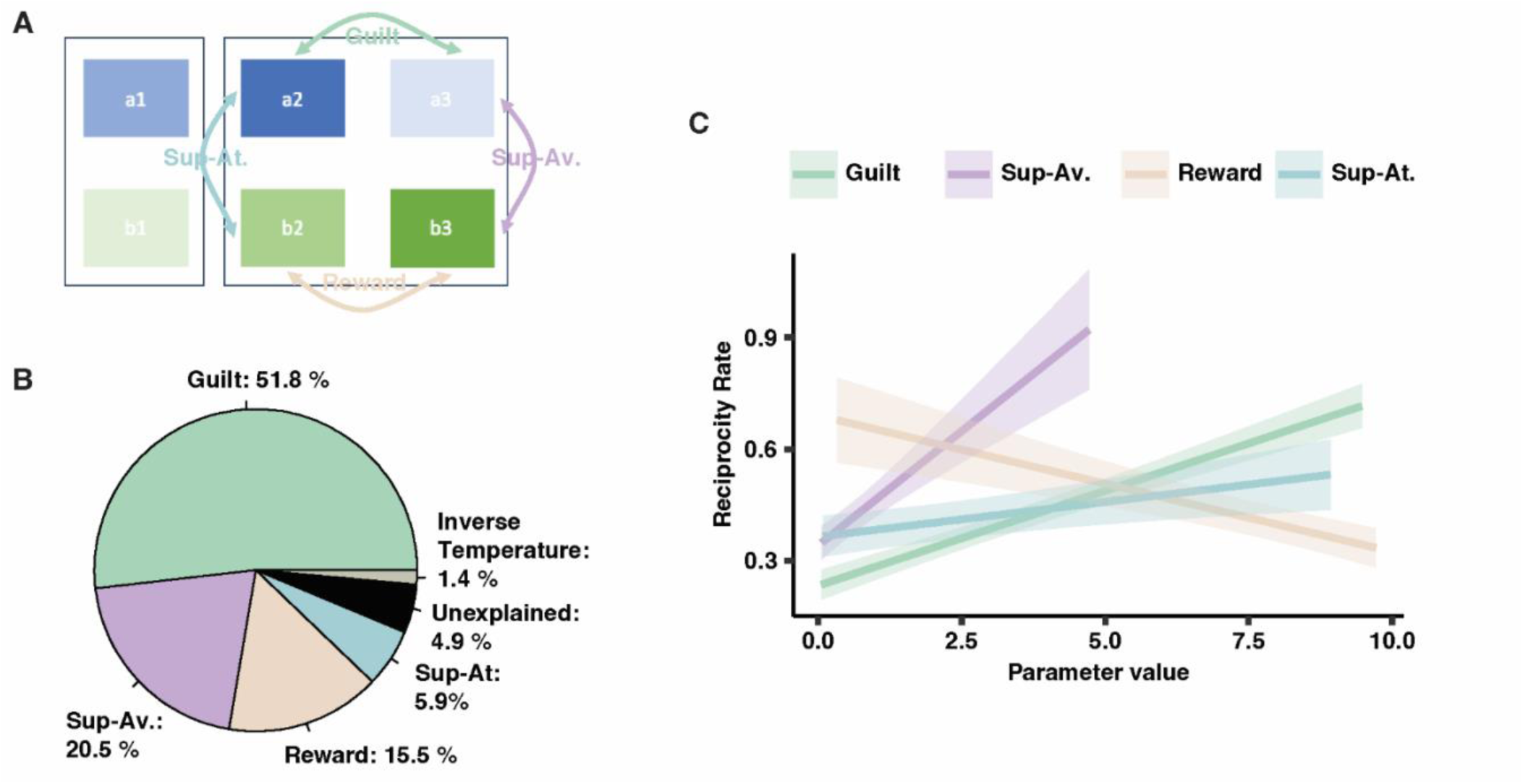
Components in the winning model and their association with reciprocity. **(A) Schematic of winning model.** Schematic illustration of the payoff structure in the binary trust game with components of the winning model: guilt aversion, superiority aversion, reward sensitivity, and superiority attraction. The blue-filled areas (a2>a1>a3) were related to the payoff structure of the trustor (partner) and the green-filled areas (b3>b2>b1) for the trustee (participant) in both gain and loss frames (Note that deeper color signifies higher value of gain or lower value of lose). The objective size of guilt aversion was quantified as the difference in the payoff structure between a2 and a3; superiority aversion between b3 and a3; reward between b3 and b2; and superiority attraction between b2 and a2. **(B) Variance explained by components.** The four components explained more than 93% of the variance of the reciprocity rate. Inverse temperature (1.4%) controls the trade-off between randomness and determinism in decision-making processes and there is a 4.9% variance unexplained by the model. **(C) Association between parameters of components and reciprocity rate.** Higher subjective sensitivity to guilt aversion, superiority aversion, and superiority attraction were associated with increased reciprocity rates, whereas higher sensitivity to reward was associated with lower reciprocity rates. Guilt: guilt aversion; Sup-Av.: superiority aversion; Sup-At.: superiority attraction; Reward: reward sensitivity.

**Table 1.** Model comparison. The winning model M4 outperformed all other candidate models, as indicated by LOOIC and WAIC. **M4** included the components of reward sensitivity (***β_R_***), guilt aversion ***β_G_***, superiority aversion ***β_SAv_***, superiority attraction ***β_SAt_*** and Inverse tempareture (***λ***) for four conditions (Anxiety [*high, low*] × Frame [*gain, loss*]). *ΔLOOIC, leave-one-out information criterion relative to the winning model; ΔWAIC, widely applicable information criterion relative to the winning model*.

| *Models* | *Parameters* | *Number of*<br>*parameter components* | *$\Delta LOOIC$* | *$\Delta WAIC$* |
| --- | --- | --- | --- | --- |
| **M4** | $\beta_R, \beta_G, \beta_{SAv}, \beta_{SAt}, \lambda$ | **20** | **0** | **0** |
| M3 | $\beta_R, \beta_G, \beta_{SAv}, \beta_{DisAd-I}, \lambda$ | 20 | 10.9 | 8.2 |
| M5 | $\beta_R, \beta_G, \beta_{SAv}, \beta_{SAt}, \lambda$ | 20 | 11.4 | 10.5 |
| M2 | $\beta_G, \beta_{SAv}, \beta_{DisAd-I}, \lambda$ | 16 | 13.5 | 21 |
| M6 | $\beta_R, \beta_G, \beta_{SAv}, \beta_{SAt}, \lambda$ | 18 | 23.1 | 27.5 |
| M1 | $\beta_G, \beta_I, \lambda$ | 12 | 55.8 | 59.5 |
| M7 | $\beta_R, \beta_G, \beta_{SAv}, \beta_{SAt}$ | 16 | 51926.5 | 128630645 |

Model prediction from **M4** demonstrated that the true and simulated reciprocity rates were highly correlated (*rs* > 0.96, **Fig. S2**). Parameter recovery for **M4** also indicated successful recovery of all parameters from **M4** (guilt aversion: *rs* > 0.90; superiority aversion: *rs* > 0.69; reward: *rs* > 0.64; superiority attraction: *rs* > 0.68; inverse temperature: *rs* > 0.55; **Fig. S3**).

### 2.2. Validation of Winning Model through Eye-Tracking Data

To validate the winning model, the relationship between the estimated parameters of the components and observed eye movements was investigated. Six areas of interest (AOIs) were defined, corresponding to the six rectangles containing payoffs in the binary trust game (**Fig. 2A**). The relative time spent on transitions between each pair of AOIs was calculated based on the fixation sequence of eye movements within each trial (see **Materials and Methods**). As a result, 21 unique transitions were extracted and the proportion of fixation time (normalized with the total reaction time for each trial) on each transition was calculated.

Validating our winning model (**M4**), individuals spent significantly more time on the transitions related to its four identified components compared to all other transitions (**Fig. 3A**). The Linear Mixed Model (LMM) (**Tab. 2**) indicated that greater subjective sensitivity to specific components was associated with more time spent on transitions related to those components (**Fig. 3B**). Furthermore, resembling the relationship between model parameter and reciprocity rate (**Fig. 2C**), increased fixation time on transitions involving guilt aversion, superiority aversion, and superiority attraction correlated with a higher rate of reciprocity, while increased fixation time on reward transitions was linked to a lower rate of reciprocity (**Fig. 3C**, **Tab. 2**). These eye-movement results confirmed that the parameter estimated by the winning model reflects the underlying psychological components of reciprocity.

**Figure 3.**
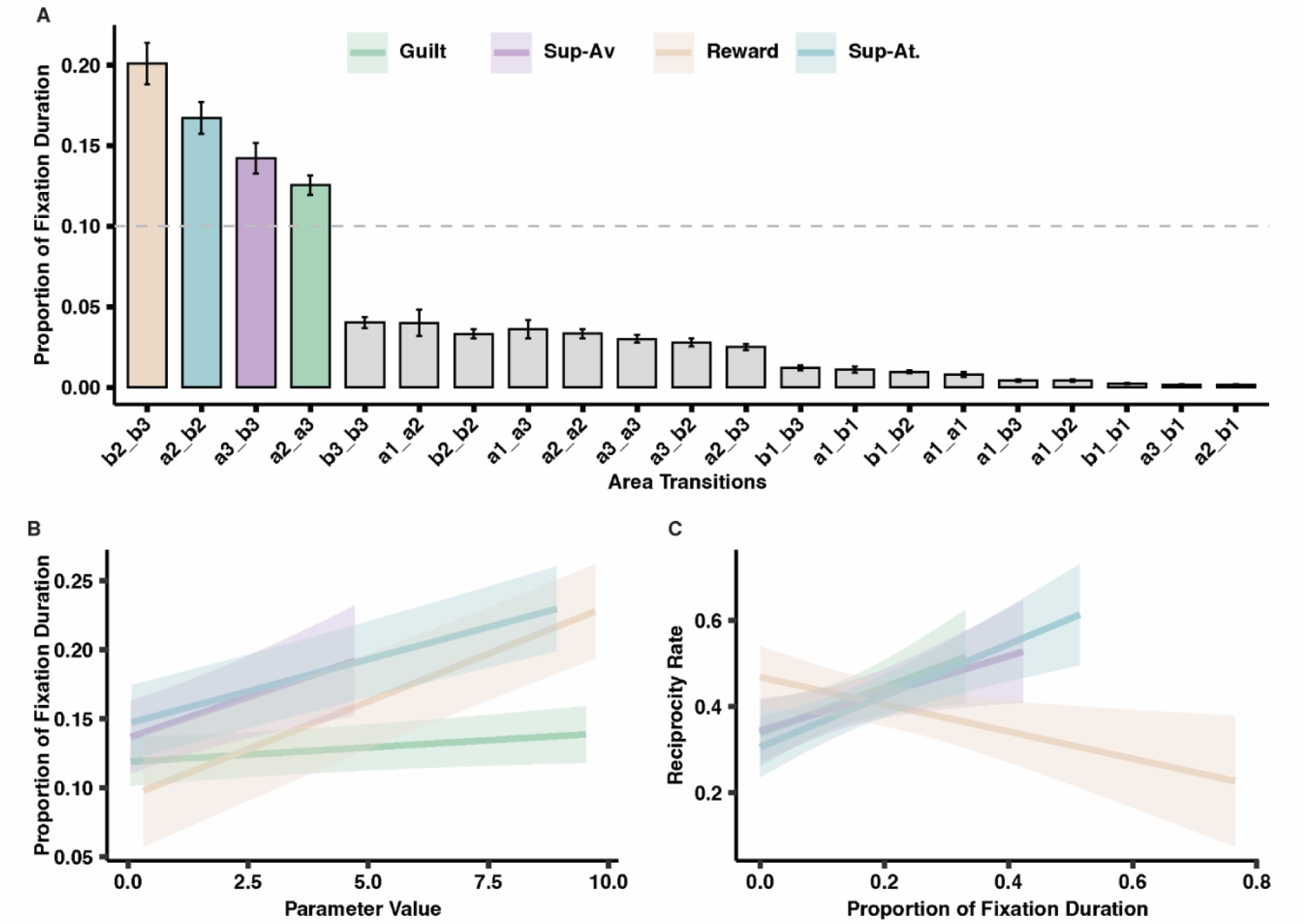
The fixation duration of the transitions and their associations with component parameters and reciprocity rate. **(A) Proportion of fixation duration on each transition (Mean ± standard error of mean, SE).** The transitions related to the four components (guilt aversion [a2_a3], superiority aversion [a3_b3], superiority attraction [a2_b2], and reward [b2_b3]) dominated the proportion of fixation duration when making reciprocity decisions. Dash line represent a 10% proportion of fixation duration. **(B) Association between parameter value of components and fixation duration.** Higher sensitivity to the components was associated with an increased proportion of fixation duration on related transitions. **(C) Association between fixation duration and reciprocity rate.** An increased proportion of duration on the transitions of guilt aversion, superiority aversion, and superiority attraction predicted a higher rate of reciprocity, while an increased proportion of duration on the transitions of reward predicts a lower rate of reciprocity. Guilt: guilt aversion; Sup-Av.: superiority aversion; Sup-At.: superiority attraction; Reward: reward sensitivity.

**Table 2.**
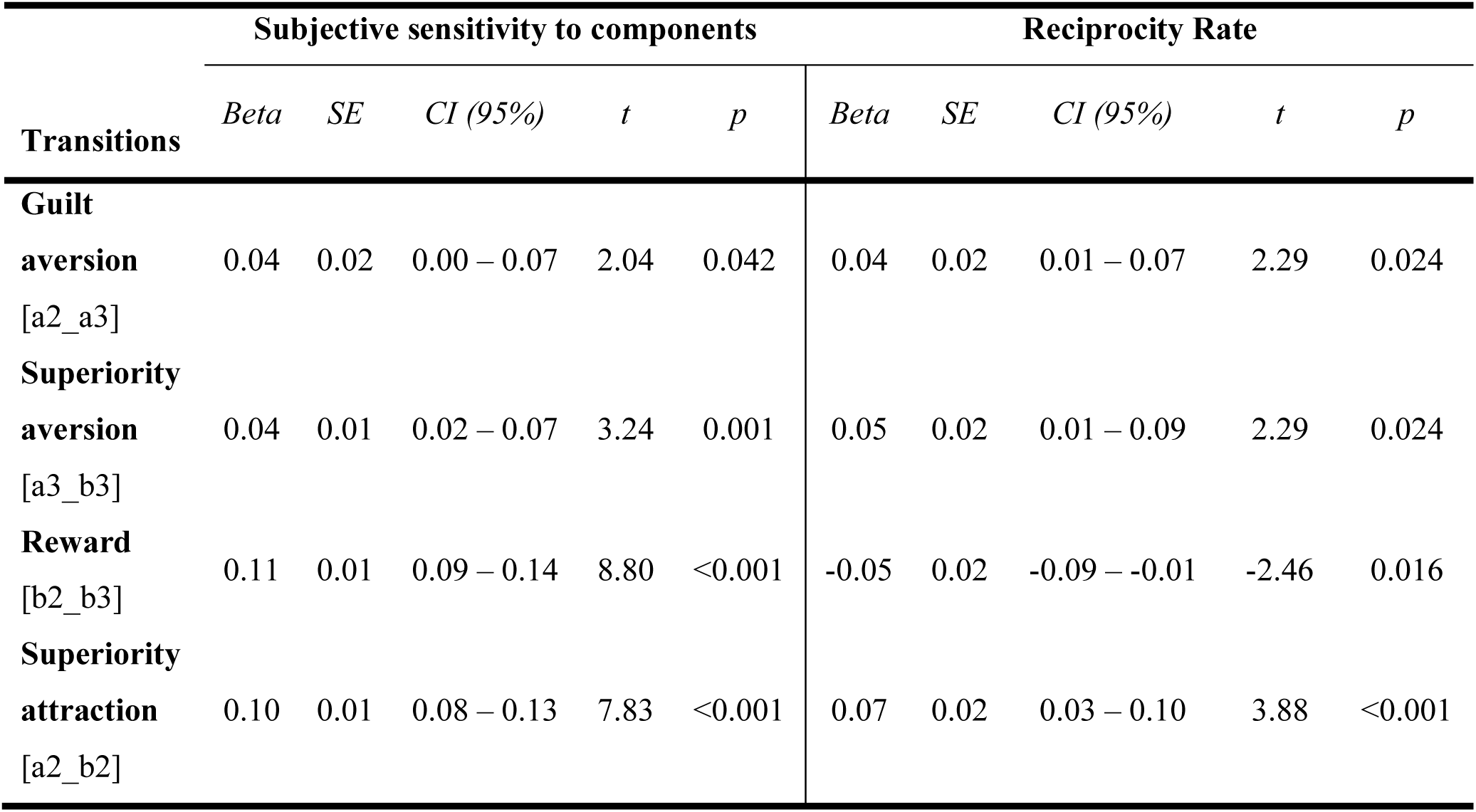
Association between fixation duration on components and subjective sensitivity to corresponding components and reciprocity rate. Left columns: An increased proportion of fixation duration of components was associated with higher subjective sensitivity to the corresponding components. **Right columns:** increased proportion of duration on the transitions of guilt aversion [a2_a3], superiority aversion [a3_b3], and superiority attraction [a2_b2] predicted a higher rate of reciprocity, while an increased proportion of duration on the transitions of reward [b2_b3] predicts a lower rate of reciprocity. SE: standard error of mean; CI (95%): 95% confidence interval.

### 2.3. Impact of Trait Anxiety on Core and Periphery of Reciprocity

The LMM on the reciprocity rate revealed a significant main effect of Anxiety, χ²(1) = 4.37, *p* = 0.036, 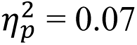 (**Fig. 4**), where individuals with high trait anxiety showed a lower reciprocity rate compared to those with low trait anxiety. The main effect of Frame was marginally significant, χ²(1) = 3.20, *p* = 0.074, 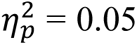, where individuals under the Gain Frame reciprocated more than those under the Loss Frame. The Anxiety × Frame interaction effect was not significant, χ²(1) = 0.003, *p* = 0.954. The presence of the main effect of Anxiety and the absence of Anxiety × Frame interaction effect suggest that trait anxiety attenuated reciprocity regardless of contexts.

**Figure 4.**
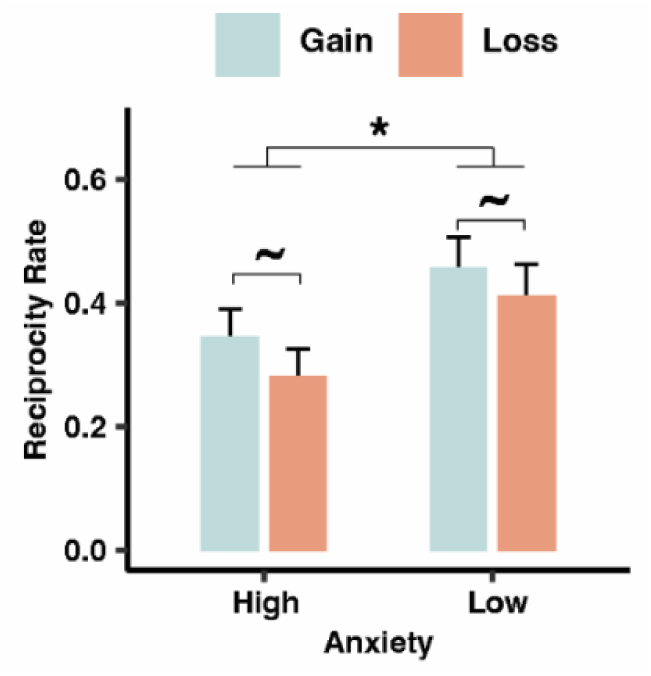
The impact of anxiety on reciprocity rate (Mean ± SE). High Anxiety group exhibited lower reciprocity rates than Low Anxiety group. Further, individuals under a Loss Frame showed a trend toward a lower reciprocity rate compared to those under a Gain Frame. **\***: *p* < 0.05, **∼**: *p* < 0.1.

The LMM on the reaction time revealed no significant main effect of Anxiety, χ²(1) = 0.21, p = 0.646, nor a significant Anxiety x Frame interaction, χ²(1) = 1.49, p = 0.221, but a marginal main effect of Frame, χ²(1) = 3.06, p = 0.085, with slower decisions under the Loss Frame than the Gain Frame (**Fig. S1**).

#### 2.3.1. Computational mechanism underlying trait anxiety’s impact on core and periphery of reciprocity

The parameter *β*_*G*_, *β*_*SAv*_, *β*_*R*_ and *β*_*SAt*_ in **M4** represent the participant’s sensitivity to guilt aversion, superiority aversion, reward, and superiority attraction, respectively.

##### Guilt aversion

The LMM on the guilt aversion parameters revealed no significant main effect of Frame, χ²(1) = 0.22, p = 0.638, but a significant main effect of Anxiety, χ²(1) = 4.23, *p* = 0.039, 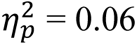, where the High Anxiety group showed lower guilt aversion compared to the Low Anxiety group (**Fig. 5A**). The main effect of Anxiety is significant and the interaction effect of Anxiety × Frame was not, χ²(1) = 0.09, *p* = 0.759, indicating that trait anxiety attenuated guilt aversion regardless of contexts.

**Figure 5.**
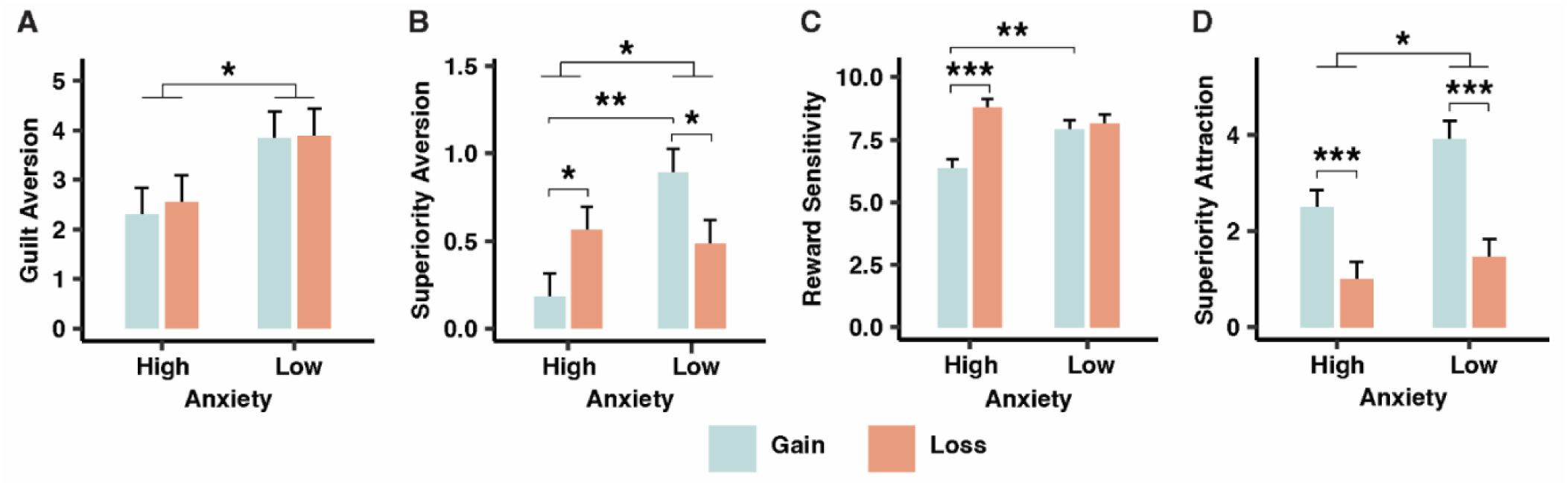
Impact of trait anxiety on the psychological components of reciprocity (Mean ± SE). (A) Guilt Aversion. Trait anxiety attenuated guilt aversion regardless of context. **(B) Superiority aversion.** Trait anxiety altered contextual perception, with high and low trait anxiety individuals showing reversed contextual effects. In addition, high trait anxiety individuals demonstrated a significant contextual effect, whereas low trait anxiety individuals showed no contextual effect. **(C) Reward sensitivity.** Trait anxiety altered the contextual perception of reward sensitivity, with high trait anxiety individuals displaying a contextual effect, while those with low trait anxiety showed no contextual effect. **(D) Superiority attraction**. Trait anxiety reduced superiority attraction regardless of context, and the loss frame decreased superiority attraction compared to the gain frame. **\*\*\***: *p* < 0.001; \***\***: *p* < 0.01; **\***: *p* < 0.05.

##### Superiority aversion

The LMM on superiority aversion parameters showed no significant main effect of Frame, χ²(1) = 0.01, p = 0.921, but a significant main effect of Anxiety, χ²(1) = 4.53, *p* = 0.033, 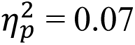, where the High Anxiety group had a lower superiority aversion compared to the Low Anxiety group (**Fig. 5 B**). Further, the interaction effect of Anxiety × Frame was significant, χ²(1) = 11.65, *p* < 0.001, 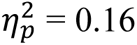. Planned follow-up post-hoc analyses revealed that Low Anxiety group showed lower superiority aversion in the Loss than in the Gain Frame, β = 0.41, SE = 0.17, t = 2.46, *p* = 0.017, but a reverse pattern was observed for the High Anxiety group, β = −0.38, SE = 0.16, t = −2.36, *p* = 0.021. The presence of a reversed contextual effect on superiority aversion between the High and the Low Anxiety group suggests that trait anxiety altered the contextual perception of superiority aversion. Moreover, High Anxiety group exhibited lower superiority aversion than those in the Low Anxiety group under the Gain Frame, β = 0.71, SE = 0.19, t = 3.78, *p* = 0.003, but not Loss Frame, β = 0.08, SE = 0.19, t = 0.42, *p* = 0.675. The presence of anxiety effect on superiority aversion in the Gain Frame but absence in the Loss Frame suggests that the effect of anxiety on superiority aversion depended on contexts.

##### Reward sensitivity

The LMM on reward sensitivity parameters revealed no significant main effect of Anxiety, χ²(1) = 1.37, p = 0.242, but a significant main effect of Frame, χ²(1) = 20.77, *p* < 0.001, 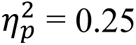, where individuals under the Loss Frame showed significantly higher reward sensitivity compared to the Gain Frame (**Fig. 5C**). Further, the interaction effect of Anxiety x Frame was significant, χ²(1) = 14.03, *p* < .001, 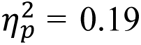. Planned follow-up post-hoc analyses revealed that individuals with high anxiety showed greater reward sensitivity in the Loss than in Gain Frame, β = −2.43, SE = 0.41, t = −5.92, *p* < 0.001. However, this pattern disappeared in the Low Anxiety group, β = −0.24, SE = 0.42, t = −0.57, *p* = 0.571. The presence of contextual effect on reward sensitivity in the High but absence in the Low Anxiety group suggests that trait anxiety altered the contextual perception of reward sensitivity. Moreover, High Anxiety group exhibited lower reward sensitivity than those in the Low Anxiety group under the Gain Frame, β = −1.55, SE = 0.49, t = −3.19, *p* = 0.002, but not Loss Frame, β = 0.64, SE = 0.49, t = 1.32, *p* = 0.189. The presence of anxiety effect on reward sensitivity in the Gain Frame but absence in the Loss Frame suggests that the effect of anxiety on reward sensitivity depended on contexts.

##### Superiority attraction

The LMM on superiority attraction parameters indicated a significant main effect of Anxiety, χ²(1) = 5.34, p = 0.021, 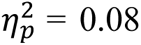, demonstrating the High Anxiety group showed lower superiority attraction than the Low Anxiety group (**Fig. 5D**). Further, a significant main effect of Frame was observed, χ²(1) = 35.40, *p* < 0.001, 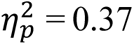, with a higher superiority attraction in the Gain compared to the Loss Frame. However, the interaction effect of Anxiety x Frame was not significant, χ²(1) = 2.11, p = 0.147, suggesting that trait anxiety attenuated superiority attraction regardless of contexts.

#### 2.3.2. Brain mechanism underlying trait anxiety’s impact on core and periphery of reciprocity

Given that guilt aversion and superiority attraction are psychological components affected by trait anxiety in the core of reciprocity (i.e., Anxiety effect regardless of contexts), and superiority aversion and reward in the periphery (i.e., Anxiety effect depended on contexts), the brain mechanisms underlying the impact of trait anxiety on core and periphery were investigated. First, ERP components of P2, N2, and LPP were examined, differentiating the impact of trait anxiety on the core and peripheral processes.

##### P2 component

The LMM analysis on the P2 amplitude showed a significant main effect of Anxiety, χ²(1) = 4.28, *p* = 0.039, 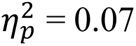, where the High Anxiety group exhibited a lower P2 amplitude than the Low Anxiety group (**Fig. 6A, B**). No main effect of Frame, χ²(1) = 2.50, *p* = 0.114, nor an interaction effect of Anxiety × Frame was found, χ²(1) = 0.05, *p* = 0.829. The presence of the Anxiety main effect and absence of Anxiety × Frame interaction effect suggests that trait anxiety decreased P2 amplitude regardless of contexts.

**Figure 6.**
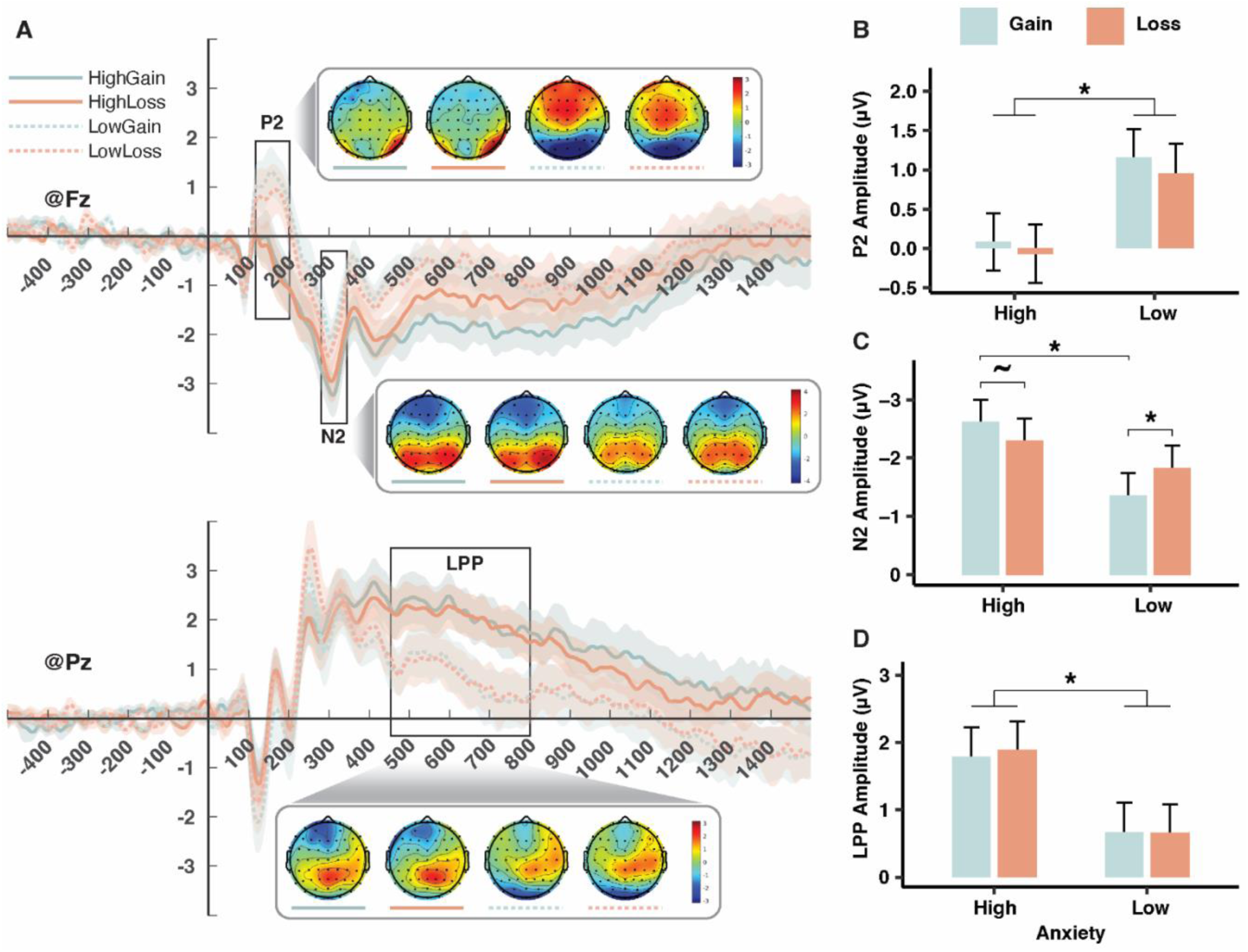
Condition comparisons of event-related potentials on reciprocity decision (Mean ± SE). (A) Grand average potential waveforms. Grand average potential waveforms for the four conditions (Anxiety × Frame) were measured at the Fz electrode and Pz. **(B) Comparison of P2 amplitudes.** P2 amplitudes (measured at F1, Fz, F2) were significantly lower in individuals with high trait anxiety compared to those with low trait anxiety regardless of frame. **(C) Comparison of N2 amplitudes**. N2 amplitudes (measured at F1, Fz, F2) showed a reversed contextual effect for individuals with high and low trait anxiety. **(D) Comparison of LPP amplitudes.** LPP amplitude (measured at P1, Pz, P2) was significantly lower in individuals with high trait anxiety than those with low trait anxiety regardless of frame. **\***: p < 0.05, **∼**: p < 0.1.

##### N2 component

The LMM analysis on the N2 amplitude demonstrated no significant main effect of Frame, χ²(1) = 0.32, p = 0.573, nor a main effect of Anxiety, χ²(1) = 2.67, p = 0.102, where the High Anxiety group exhibited a lower N2 amplitude than the Low Anxiety group (**Fig. 6A, C**). Further, a significant Anxiety × Frame interaction effect was observed, χ²(1) = 9.46, *p* < 0.001, 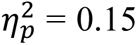. Planned follow-up post-hoc analyses indicated that the Low Anxiety group showed a greater N2 amplitude under the Loss compared to the Gain Frame, β = −0.46, SE = 0.18, z = −2.56, *p* = 0.011. However, the High Anxiety group showed a reversed pattern, demonstrating a marginally significantly smaller N2 amplitude under the Loss compared to the Gain Frame, β = 0.32, SE = 0.18, z = 1.79, *p* = 0.074. The presence of a reversed contextual effect on N2 amplitude between the High and the Low Anxiety group suggests that trait anxiety altered the contextual effect of N2 amplitude. Moreover, High Anxiety group exhibited a smaller N2 amplitude than Low Anxiety group under the Gain Frame, β = 1.26, SE = 0.54, z = 2.32, *p* = 0.020, but not Loss Frame, β = 0.47, SE = 0.55, z = 0.87, *p* = 0.387. The presence of anxiety effect on N2 amplitude in the Gain Frame but absence in the Loss Frame suggests that the effect of trait anxiety on N2 amplitude depended on contexts.

##### LPP component

The LMM analysis on the LPP amplitude showed a significant main effect of Anxiety, χ²(1) = 4.19, *p* = 0.041, 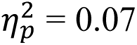, where the High anxiety group exhibited a higher LPP amplitude than the Low Anxiety group (**Fig. 6A, D**). However, no significant main effect of Frame, χ²(1) = 0.12, *p* = 0.716, nor interaction effect of Anxiety × Frame were found, χ²(1) = 0.23, *p* = 0.633. The presence of Anxiety main effect and absence of Anxiety × Frame interaction effect suggests that trait anxiety decreased LPP amplitude regardless of contexts.

Second, mediation analyses were employed to investigate how trait anxiety impacted these ERP components and, subsequently, the psychological components underlying reciprocity. Since trait anxiety affected P2 and LPP responses similarly to guilt aversion and superiority attraction regarding the core of reciprocity, serial mediation analysis was used to test if trait anxiety impacts these psychological components through changes in P2 and LPP amplitudes.

The serial mediation analysis revealed that higher trait anxiety was associated with a lower P2 amplitude and a higher LPP amplitude, leading to reduced guilt aversion (indirect effect [a*b*c] = −0.017, SE = 0.003, z = −6.34, *p* < 0.001, 95% CI [-0.022, −0.012]) (**Fig. 7A**). Further, higher trait anxiety attenuated guilt aversion through increased LPP amplitude alone (indirect effect [a2*c] = −0.090, SE = 0.015, z = - 6.15, *p* < 0.001, 95% CI [-0.121, −0.063]). Moreover, trait anxiety did not significantly modulate guilt aversion through P2 amplitude alone (indirect effect [a*b2] = 0.019, SE = 0.015, z = 1.23, *p* = 0.220, 95% CI [-0.012, 0.048]). Those results demonstrated that higher trait anxiety led to lower guilt aversion by increased LPP amplitude alone or by reducing P2 amplitude, which then translated into increased LPP amplitude.

**Figure 7.**
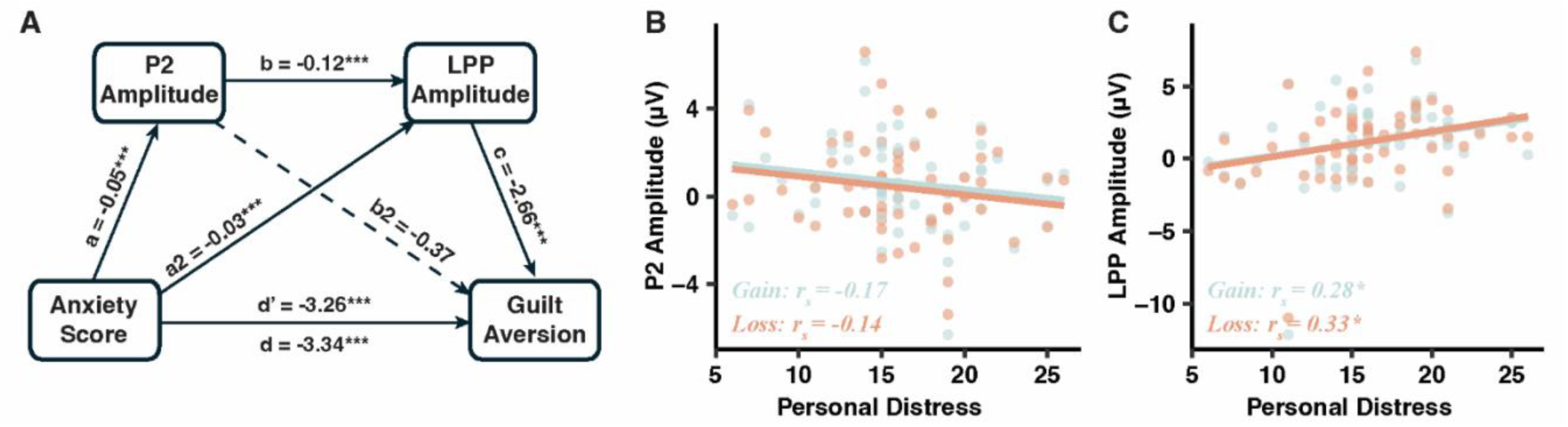
P2 and LPP mediated trait anxiety ef fects on guilt aversion and their association with personal distress. **(A)** The attenuating effect of trait anxiety on guilt aversion occurred partially through decreasing P2 amplitude and increasing LPP amplitude. While **(B)** Personal distress is not related to P2 amplitude, **(C)** but positively related to LPP amplitude. **\*\*\***: *p* < 0.001; \***\***: *p* < 0.01; **\***: *p* < 0.05.

However, the serial mediation analysis showed no significant indirect effect of trait anxiety on superiority aversion through both P2 and LPP amplitude (indirect effect [a*b*c] = 0.001, SE = 0.001, z = 1.01, *p* = 0.311, 95% CI [-0.000, 0.002]), nor effects through P2 amplitude alone (indirect effect [a*b2] = 0.009, SE = 0.006, z = 1.40, *p* = 0.163, 95% CI [-0.003, 0.022]) or LPP amplitude alone (indirect effect [a2*c] = 0.003, SE = 0.003, z = 1.02, *p* = 0.309, 95% CI [-0.003, 0.010]). These insignificant results suggest that P2 or LPP amplitude was not involved in the processing of superiority attraction.

Third, to better interpret the ERP components, the associations between ERP amplitudes and self-report measures like IRI (empathy) and Machiavellianism (selfishness) were examined. Mann-Whitney U tests showed that the High Anxiety group exhibited higher scores on Machiavellianism and IRI personal distress compared to the Low Anxiety group (**Tab. 3**). Further, Spearman correlation analyses indicated that Machiavellianism was not correlated with P2 amplitude (Gain: *r_s_* = −0.14, *p* = 0.294; Loss: *r_s_* = - 0.074, *p* = 0.576) nor LPP amplitude (Gain: *r_s_* = 0.17, *p* = 0.186; Loss: *r_s_* = 0.12, *p* = 0.373). In contrast, personal distress was not correlated with P2 amplitude (Gain frame: *r_s_* = −0.17, *p* = 0.196; Loss frame: *r_s_* = −0.14, *p* = 0.302) (**Fig. 7B**), but positively associated with LPP amplitude (Gain frame: *r_s_* = 0.28, *p* = 0.027; Loss frame: *r_s_* = 0.33, *p* = 0.011 (**Fig. 7C**). These results suggest that LPP amplitude may reflect emotional regulation mechanisms for negative emotions, especially in individuals with high trait anxiety. Individuals who experienced higher levels of personal distress, tended to exert more effort to suppress anticipatory guilt (indicated by higher LPP amplitude) as an emotional regulation strategy to avoid further emotional burden during social interactions.

**Table 3.** Psychometric measures (Mean ± SE). Individuals with high trait anxiety group exhibited higher scores on Machiavellianism and IRI personal distress compared to those with low anxiety. SE: standard error of mean; *U*: Mann-Whitney *U* value; *p: p-value*.

|  | High trait anxiety | Low trait anxiety | *U* | *p* |
| --- | --- | --- | --- | --- |
| Machiavellianism | 79.47±2.35 | 71.87±1.74 | 604 | 0.023 |
| Interpersonal Reactivity Index |  |  |  |  |
| Perspective-taking | 18.03±0.79 | 19.35±0.63 | 357.5 | 0.172 |
| Fantasy | 17.72±0.63 | 17.77±0.85 | 457.5 | 0.917 |
| Empathic concern | 19.19±0.64 | 19.39±0.57 | 428.5 | 0.755 |
| Personal distress | 17.91±0.63 | 14.71±0.83 | 636.5 | 0.006 |

#### 2.3.3. Brain mechanism of trait anxiety on contextual effect

Since N2 reflects cognitive control, which is essential for contextual perception and influenced by trait anxiety on the periphery, the relationship between N2 amplitude, superiority aversion, and reward sensitivity was tested. The contextual effect of superiority aversion and reward sensitivity was calculated as the difference between loss and gain contexts. The impact of trait anxiety on the contextual effect of N2 amplitude, superiority aversion, and reward was analyzed.

Mann-Whitney U tests showed that Anxiety significantly altered the contextual effect on superiority aversion (U = 128, *p* < 0.001 (**Fig. 8A**) and N2 amplitude (U = 316, *p* = 0.048) (**Fig. 8B**) in a similar manner. Spearman correlation showed that changes in the contextual effect on superiority aversion positively correspond with changes in N2 amplitude (*r_s_* = 0.38, *p* = 0.003) (**Fig. 8C**). Mann-Whitney U tests showed that Anxiety significantly altered the contextual effect on reward sensitivity (U = 227, *p* < 0.001). However, Spearman correlation indicated no significant relationship between the contextual effect of reward sensitivity and N2 amplitude (*r_s_* = 0.15, *p* = 0.245). These results suggest that N2 amplitude potentially regulates the contextual perception of superiority aversion.

**Figure 8.**
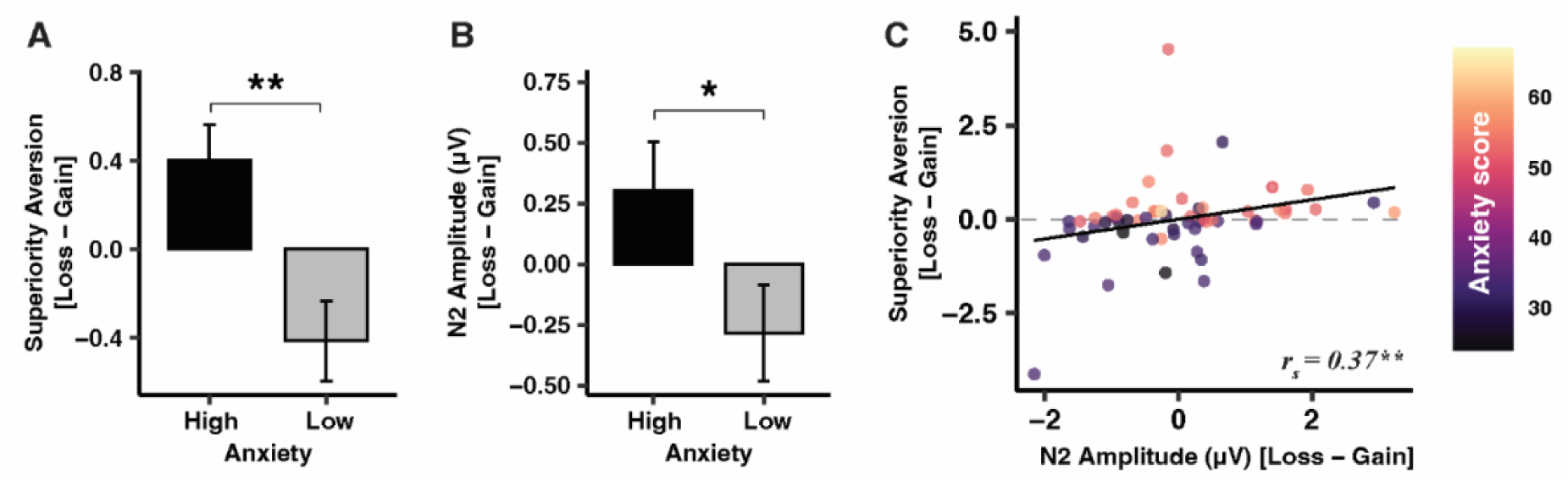
Alteration of trait anxiety on contextual perception of superiority aversion and N2 mechanism. **(A) Alteration of trait anxiety on the contextual perception of superiority aversion.** Trait anxiety significantly altered the contextual effect of superiority aversion. **(B) Alteration of trait anxiety on the contextual effect of N2 amplitude.** Similarly, trait anxiety affected N2 amplitude in a parallel pattern with superiority aversion. **(C) Association between the contextual effect of N2 amplitude and those of superiority aversion**. The contextual effect of N2 amplitude was positively correlated with the contextual effect of superiority aversion. \***\***: *p* < 0.01; **\***: *p* < 0.05.

## 3. Discussion

Reciprocity is a complex social behavior influenced by both reciprocity propensity (core) and contextual perception (periphery) [4,5]; however, their underlying neurocomputational mechanism remain unknown. In this study, we employed computational modeling, eye-tracking, and ERP within an economic binary trust game under gain and loss frame to examine how trait anxiety affects the core and periphery of reciprocity and its underlying psychological components. We identified four psychological components— validated by eye-tracking analyses—underlying reciprocity decisions: reward, guilt aversion, superiority aversion, and superiority attraction. For the core of reciprocity decisions, we found that trait anxiety reduced reciprocity at the behavioral level and decreased guilt aversion and superiority attraction at the psychological level across different framed contexts. At the neural level, we found that trait anxiety attenuated guilt aversion by decreasing P2 amplitude related to selective attention and increasing LPP amplitude linked to effortful emotion regulation. For the periphery of reciprocity decisions, we showed that trait anxiety altered the contextual perception of reward and superiority aversion at the psychological level, but did not affect reciprocity at the behavioral level. In particular, trait anxiety altered the contextual perception of superiority aversion which involved the N2 cognitive control mechanism at the neural level. Overall, our study added knowledge on the neurocomputational mechanisms of how trait anxiety impacts the core and periphery of reciprocity decisions.

### 3.1. Computational Modeling of Reciprocity Decisions

Using computational modeling, we first analyzed the psychological components of how individuals make reciprocity decisions. In line with our first hypothesis, extending previous findings [8,9,17], our best model identified four psychological components of reciprocity decisions: reward, guilt aversion, superiority aversion, and superiority attraction. Individuals who prioritize reward tended to reciprocate less, while those who emphasize guilt aversion, superiority aversion, and superiority attraction were more likely to reciprocate. Our model indicated that superiority aversion occurs when evaluating the betrayal option, while superiority attraction appears when evaluating the reciprocity option in the binary game. Although the reward in the reciprocity option is less appealing compared to the betrayal option, individuals are more likely to reciprocate due to the higher relative payoff, supporting the notion that superiority aversion is condition-dependent rather than always present [10,14–16].

Next, we validated the model’s predicted psychological components for reciprocity decisions using eye-prediction and parameter recovery, eye-tracking results also empirically validated our model. Our results showed that the transitions of the four components—guilt aversion, superiority aversion, reward, and superiority attraction—dominated the top of all transitions, indicating the significant contribution of these components in reciprocity decision-making. Furthermore, higher component parameter values were correlated with increased relative fixation duration on the corresponding transitions, based on the binary trust game’s payoff structure. Moreover, resembling the relationship between the four components and reciprocity rate, we found that increased fixation duration on transitions mapping guilt aversion, superiority aversion, and superiority attraction correlated with a higher reciprocity rate, while increased fixation duration on transition mapping reward correlated with a lower reciprocity rate. Notably, these transitions involve specific comparisons and evaluations during decision-making as a trustee based on the payoff structure in the binary trust game [18–20]. For reward, it involves comparing the participant’s payoffs between reciprocity and betrayal options, whereas for guilt aversion, it involves comparing the partner’s payoffs between these options. Superiority attraction is evaluated by comparing payoffs between the participant and the partner within the reciprocity option, while superiority aversion is assessed by comparing these payoffs within the betrayal option.

### 3.2. Trait Anxiety’s Impact on the Core of Reciprocity

To explore how trait anxiety affects reciprocity, we examined the behavioral and computational mechanisms in both gain and loss contexts. We partially confirmed our third hypothesis that trait anxiety influences reciprocity both at the behavioral and psychological levels regardless of context. At the behavioral level, consistent with previous studies [23–25], our results indicated that individuals with high trait anxiety exhibited lower reciprocity than those with lower trait anxiety. At the psychological level, our results demonstrated that trait anxiety attenuated guilt aversion as well as superiority attraction independently of context. These effects align with findings in individuals with OCPD, often comorbid with anxiety disorders [17]. Anxious individuals tend to use avoidance strategies in social interactions [26–28], likely reducing guilt aversion by limiting affective processing and diminishing a sense of moral obligation [17,57].

Contrary to our hypothesis, we failed to observe an impact of trait anxiety on superiority aversion as a core psychological component of reciprocity. While anxiety had a main effect on superiority aversion, the interaction between trait anxiety and frame showed that this effect depends on the context. Specifically, trait anxiety reduced reciprocity in a gain frame but had no effect in a loss frame, suggesting it impacts the periphery of reciprocity rather than its core. Although less discussed in the literature, our results showed that trait anxiety reduces sensitivity to superiority attraction when evaluating reciprocity. Due to the smaller payoff difference in the reciprocity option compared to the betrayal option, individuals with high trait anxiety might overlook minor payoff differences because of impaired attention and increased distractibility [63,64]. Alternatively, their cognitive bias toward negative outcomes might have reduced their sensitivity to positive incentives [65] like superiority attraction in reciprocity decisions.

Building on our computational findings, we investigated the neural mechanisms of how trait anxiety affects reciprocity. As hypothesized, our study revealed that higher trait anxiety is associated with lower P2 and higher LPP amplitudes, regardless of context. The P2 component is linked to selective attentional allocation [37–40], whereas the LPP component is associated with an effortful cognitive regulation over emotion [50–53], particularly during the resolution of moral conflicts [54–56]. These results suggest that individuals with high trait anxiety may allocate fewer attentional resources to evaluating moral conflicts or exert more effort to resolve or disengage from the dilemmas in reciprocity decisions.

Our mediation analysis further supports this interpretation, indicating that trait anxiety attenuated guilt aversion through diminished P2 and elevated LPP amplitudes. To further elucidate the role of these ERP components, we examined relationships between self-report measures (Machiavellianism and IRI) and P2 as well as LPP amplitudes. Both Machiavellianism and IRI personal distress were higher in individuals with high trait anxiety compared to those with low trait anxiety. Further, individuals with higher personal distress exhibited higher LPP amplitudes in both gain and loss contexts. However, no significant relationships were found between personal distress and P2 amplitudes, Machiavellianism and P2 amplitudes, or Machiavellianism and LPP amplitudes. These findings suggest that LPP reflects effortful regulation and disengagement from negative emotions [50–53], supporting the idea that individuals with higher trait anxiety tend to adopt avoidance strategies [26–28]. Overall, anxiety’s decreasing effect on P2 amplitudes indicates fewer attentional resources allocated to assessing guilt [37–40], while anxiety’s increasing effect on LPP amplitudes suggests effortful emotion regulation and disengagement from the anticipatory guilt in reciprocity decisions [54–56].

### 3.3. Trait Anxiety’s Impact on the Periphery of Reciprocity Decisions

To elucidate the trait anxiety’s impact on the periphery of reciprocity decisions, we further investigated the behavioral and computational mechanism underlying how trait anxiety alters the contextual perception between gain and loss context. Partially supporting our fourth hypothesis, trait anxiety did not affect the behavioral aspects of reciprocity but did alter its psychological components. Our results indicated that individuals with low trait anxiety showed no contextual effect on reward, while those with high trait anxiety were more sensitive to reward under a loss than a gain frame. This finding aligns with previous evidence [32–34] showing that individuals with high trait anxiety are more susceptible to contextual effects on reward. Our results further showed that trait anxiety alters the contextual effect on superiority aversion. While individuals with high trait anxiety generally exhibited lower superiority aversion, context modulated this response: a loss frame reduced superiority aversion in those with low trait anxiety but enhanced it in those with high trait anxiety. This indicates distinct context-dependent processing mechanisms between individuals with low and high trait anxiety.

Anxiety’s effect on contextual perception may stem from how individuals with varying trait anxiety levels respond to different contexts. Individuals with high anxiety tend to rely more on heuristic decision-making than those with low anxiety [33]. Further, loss frames are heuristically perceived as more harmful to others [7,36] and more threatening to one’s own reward [34] compared to gain frames. Therefore, loss framing prompts individuals with high trait anxiety to increase superiority aversion and reward sensitivity, possibly driven by the “do-no-harm” principle [7] and self-protective strategies [66]. For example, even with the same payoff structure, making a partner lose more than the decision-maker is seen as more harmful than making the partner gain less. Similarly, losing more of their own payoff is perceived as more harmful than gaining less. The dual increase in superiority aversion and reward sensitivity in high trait anxiety individuals reflects the complex reality of decision-making with conflicting effects of the latent psychological components. This may potentially explain the lack of a significant contextual effect on behavioral reciprocity in high-anxiety individuals, as the effect of these latent psychological components might cancel each other out. In contrast, low anxiety individuals showed higher superiority aversion under the gain framing but lower under loss framing, suggesting a shift toward self-protection over other-regarding behavior.

Looking into the neural mechanism underlying trait anxiety’s impact on the periphery of reciprocity, consistent with our fourth hypothesis, our findings indicate that trait anxiety reverses the pattern of N2 amplitude in line with superiority aversion. When moving from a gain to a loss context, individuals with low trait anxiety showed decreased superiority aversion and heightened N2 amplitude, while those with high trait anxiety exhibited increased superiority aversion and attenuated N2 amplitude. The context’s influence on superiority aversion was linked to its effect on N2 amplitude. The N2 component, which emerges during decision-making involving conflict and typically indicates higher cognitive control or more effortful response inhibition [67,68], is also linked to the framing effect [69]. Our results suggest several key insights: Superiority aversion appears to be an instinctual response, with reducing it requiring cognitive control and enhanced N2 effort, while increasing it involves less cognitive effort. Further, the influence of framing on superiority aversion is likely modulated by N2 cognitive control, i.e., that perception differences in superiority aversion due to framing are shaped by the degree of cognitive control exerted during decision-making. Finally, trait anxiety likely alters how framing affects superiority aversion through mechanisms that regulate cognitive control.

### 3.4. Limitation and Future Direction

Several limitations should be noted in this study. Firstly, while our model identified four components influencing reciprocity in parallel, individuals may evaluate decisions hierarchically, prioritizing some components initially and others later. Future research should model the hierarchical structure of these components [70]. Secondly, our participants were healthy college students, not clinical patients with anxiety disorders, so generalizing to clinical populations should be done cautiously. Future studies should validate these results in clinical populations. Nonetheless, our findings could aid in diagnosing anxiety disorders. For instance, individuals showing lower guilt aversion, lower P2 amplitudes, and higher LPP amplitudes during reciprocity decisions may be at higher risk for anxiety disorders. Thirdly, we measured trait anxiety but did not assess state anxiety during the experiment. Future studies should measure state anxiety to control for this potential confound. Finally, our study combined eye-tracking and EEG measures, but eye movements can create artifacts in EEG data. We used independent component analysis to minimize these artifacts, enhancing data reliability. Future studies should further refine these methods to improve data quality. Despite these limitations, our work provides valuable insights into how anxiety impacts reciprocity decisions through the lens of eye movement, EEG, and psychological components.

### 3.5. Conclusions

Our study investigated how trait anxiety affects the core and periphery of reciprocity. We found that trait anxiety reduces reciprocity propensity and influences latent psychological components such as guilt and superiority attraction, involving attentional allocation and emotion regulation processes. Additionally, trait anxiety alters the contextual effect of superiority aversion, involving cognitive control processes. Our findings offer insights into the neurocomputational mechanisms of trait anxiety’s impact on reciprocity and may aid in the improvement of reciprocity in individuals with anxiety disorders.

## 4. Materials and Methods

### 4.1. Participants

A total of 550 participants completed the online version of the trait subscale of the Chinese State-Trait Anxiety Inventory (STAI) [71,72]. The sample’s trait anxiety scores ranged with a mean (M) of 41.52 and a standard deviation (SD) of 9.75, with the 25th percentile at 35 and the 75th percentile at 48. Participants were classified into two groups based on their trait anxiety scores (TAS): those with TAS ≤ 35 were defined as the low trait anxiety group, and those with TAS ≥ 48 were defined as the high trait anxiety group. A target sample size of 30 participants for each trait anxiety group was determined based on prior related work [54–56]. In total, 69 participants were recruited for the study. Of the recruited participants, 34 were assigned to the low trait anxiety group (16 females; M = 31.65, SD = 2.83), and 35 were assigned to the high trait anxiety group (18 females; M = 53.57, SD = 3.12). None of the participants reported taking psychoactive medications or any history of mental disorder or brain injury. Participants who pressed the same button in 95% of the trials were considered nonresponsive to the task setting or not focused on the task, and were excluded from the analysis. As a result, 63 participants were included in the analysis, with 31 in low trait anxiety group (15 females; M = 31.65, SD = 2.85), and 32 in high trait anxiety group (16 females; M = 53.56, SD = 3.22). The study was conducted according to the Declaration of Helsinki and approved by the local Ethics Committee at Shenzhen University, China. Participants provided written informed consent prior to their involvement in the study. Compensation included a fixed attendance fee of 60 yuan (approximately $10) and a variable monetary reward based on their decisions during the game, which ranged from 40 yuan to 80 yuan (approximately $7 to $13).

### 4.2. Questionnaire

Participants completed two self-report questionnaires: the Interpersonal Reactivity Index (IRI) measuring four empathy subscales (perspective taking, fantasy, empathic concern, and personal distress) [58], and the Machiavellianism (Mach-IV) scale assessing selfish tendencies [59].

### 4.3. Experimental Procedure and Tasks

Prior to the experiment, participants underwent a comprehensive briefing session on the rules of the game. This session also included practice designed to familiarize participants with their roles and the structure of the binary trust game. Participants were informed that their partners had already participated as the first movers, and their decisions were recorded and stored in the computer system, with these decisions to be shown in the formal experiment. This setup was designed to avoid the potential confound brought by physical interaction with a partner on the measurement of reciprocity and also foster a belief among participants that they were engaging in authentic dynamics of interpersonal interaction.

During the experimental task, participants completed 240 trials in two counterbalanced sessions (Gain frame, Loss frame) to control for order effects, each with two 60-trial blocks separated by short breaks. Each trial paired participants with a new anonymous partner (icon, partially obscured name). Participants then chose to “reciprocate” or “betray” the partner’s “trust.” Immediate feedback revealed whether the partner (50% chance) initially “trusted” or “distrusted.“

The binary trust game, utilized in this study, is an interactive economic game involving two player roles: Player A and B [7]. In the present study, the participant played the role of B and the partner played the role of A. The rule for the binary game was as follows: A made a first move by choosing “distrust” (the square containing the left column) or “trust” (the combined square containing the middle and right columns). If A chose “distrust”, the system distributed the point directly to A (gaining [in the gain frame] or losing [in the loss frame] the points colored in blue) and to B (gaining or losing the points colored in green). In this case, B’s subsequent choice did not influence the point distribution. Conversely, if A chose “trust,” the distribution of points depended on B’s decision. B could either “reciprocate” by selecting the square on the left inside the “trust” rectangle or “betray” by selecting the square on the right inside the same rectangle. Based on B’s decision, both players would gain or lose points accordingly.

To mitigate the potential influence of decision spatial location on reciprocity choices, various versions of the binary trust game were employed. These adaptations involved altering the positions of the “trust” and “distrust” options, as well as the “reciprocate” and “betray” options, by switching their placements from left to right and vice versa. Consequently, four distinct versions of the game structure were applied. These variations were counterbalanced across participants in each trait anxiety group to ensure that any effects related to the positioning of choices were minimized, allowing for a more accurate assessment of decision-making behaviors without the bias of spatial location. Note that all four versions were standardized into one version (as displayed in **Fig. 2A**) for statistical analysis. The potential influence of the different versions was examined to ensure that the experimental effects were not due to version differences (**Supplementray**).

Trials were varied by changing the six payoff values. The payoffs structure in all the trials in both gain and loss frames has several features (**Fig. 2A**): **(1)** for A, a2 > a1 > a3; **(2)** for B, b3 > b2 > b1; and **(3)** while a1>b1 and a3 < b3 in all trials, a2 can be greater than, equal to, or less than b2 in different trials, with a2 > b2 in 124 trials, a2 = b2 in 6 trials, and a2 < b2 in 110 trials. Thus, rationally, to maximize the outcome, A should “trust” and expect B to “reciprocate”, securing the payoff a2, which is economically the most beneficial for A. Conversely, B should consistently choose to “betray” to attain b3, which is economically the most beneficial for B.

The loss frame was constructed from the gain frame using a modified version [7], balancing the value scale between frames. Unlike Evans & van Beest (2017), who used b3 (highest gain frame value) as reference, we used the sum of “betray” option values (a3 + b3) to construct the loss frame. The six loss frame values were calculated as the difference between each corresponding gain frame value and the reference value (e.g., 59 [14+45]) (**Fig. 1**). In the gain frame, participants started with 0 points and gained points throughout. In the loss frame, participants started with 15000 points and lost points based on decisions, ensuring equivalent final outcomes if the same strategies were used in both frames [7]. Final income was randomly determined from either the accumulated points in the gain session or the remaining points in the loss session.

### 4.4. Eye-Tracking Data Acquisition and Processing

Eye movements were recorded at 1000 Hz using an EyeLink 1000 Plus (SR Research Ltd., Ottawa, Ontario, Canada) with head-chin stabilization. Participants were seated 60 cm from a 1280×1024 pixel monitor and instructed to maintain fixation on a central cross. A 9-point calibration/validation was performed before each block. Fixations, saccades, and blinks were classified from raw data using EyeLink Data Viewer software (SR Research Ltd., Ottawa, Canada) with default settings. Subsequent analysis in R (“eyelinker” library) focused on fixations, excluding saccades and blinks. Large stimulus separation and high calibration accuracy allowed for generous AOI margins. Six rectangular AOIs (a1, a2, a3, b1, b2, b3; 164×144 pixels each) were aligned with the six values in the binary trust game (**Fig. 2A**). A central fixation cross AOI (“Center“; 105 x 85.5 pixels) was also defined, with remaining areas labeled “Undefined.” Manual examination of fixation areas for each participant and block resulted in the exclusion of four participants due to drift outside AOIs. For each trial, fixation sequences were extracted, and unique transitions between the six value AOIs (excluding “Center” and “Undefined”) were identified. These transitions, without considering direction (e.g., a2_a3 is equivalent to a3_a2), reflecting decision-maker comparisons [73], were mapped to model components like guilt aversion (a2_a3), superiority aversion (a3_b3), reward (b2_b3), and superiority attraction (a2_b2). Relative fixation time for each of the 21 possible AOI combinations was calculated by dividing fixation duration by response time, with absent transitions assigned 0. For example, in a trial with fixation sequence “center_a1_a3_a3_undefined_a2_undefined_b2_b3,” only transitions a1_a3 and b2_b3 would be relevant for decision processing.

### 4.5. EEG Data Acquisition and Processing

EEG data was collected during the binary trust game using 64 Ag/AgCl electrodes placed according to the International 10-20 system (Brain Products GmbH). Recordings were made at 1000 Hz with a 0.01-100 Hz passband, using FCz as online reference. Electrode impedance was kept below 5 kΩ. Electrooculographic (EOG) signals were also recorded to identify and remove eye movement artifacts.

EEG data were preprocessed in EEGLAB [74], an open-source toolbox in MATLAB (The MathWorks, Inc., Natick, Massachusetts, USA). Recordings were re-referenced offline to the whole brain common average and band-pass filtered (0.1-30 Hz). Ocular artifacts were identified by their contributions to EOG channels and frontal scalp distribution, and corrected using independent component analysis [74]. EEG epochs time-locked to reciprocity decision onsets were extracted using a 2,000 ms window (−500 ms to 1500 ms). Epochs with response times under 800 ms were discarded. After baseline correction using the pre-stimulus interval, epochs were visually inspected for gross movement artifacts, which were then excluded to ensure data quality.

For each participant and each trial, the mean amplitudes of P2, N2, and LPP ERP components were measured within their specific time windows and at their related electrodes. The selection of time windows and electrodes for ERP amplitude measurement was informed by previous studies and by examining the grand average ERP waveforms and scalp topographies. In particular, the P2 amplitude was measured at 110–200 ms [75–78], and the N2 amplitude was measured at 280–340 ms [41–44] after stimulus onset at frontal electrodes (F1, Fz, and F2). The LPP amplitude was measured at parietal electrodes (P1, Pz, and P2) 450–800 ms after stimulus onset [50–53]. Scalp topographies of these ERP components were computed by spline interpolation.

### 4.6. Computational Modeling of Reciprocity Decision-Making

A stage-wise model construction procedure was utilized to identify and quantify the latent psychological components influencing reciprocity behavior in our binary trust game [60–62]. This iterative approach involved sequentially refining the model based on the performance of the previous best-fitting model. Leave-One-Out Information Criterion (LOOIC) and Widely Applicable Information Criterion (WAIC) were used for model comparison to minimize overfitting, with lowest values indicating best fit to the data [60–62]. The goodness of fit was assessed using Tjur’s pseudo R² [79], with higher values indicating better model fit. Parameters were estimated using hierarchical Bayesian analysis [60–62]. The posterior inference was conducted via Markov chain Monte Carlo (MCMC) sampling, utilizing four independent chains, each with 4,000 iterations, to draw samples from the posterior distribution and ensure robust parameter estimation [61]. Seven plausible candidate models were tested in total using Rstan in R [80] for model-related procedures.

Guided by prior research [8,17], the baseline model incorporated guilt aversion, inequity aversion, and reward, positing that decisions arise from the interplay between these socio-emotional factors and potential gains. The utility function (*U*) was formulated as **Eq. 1**:

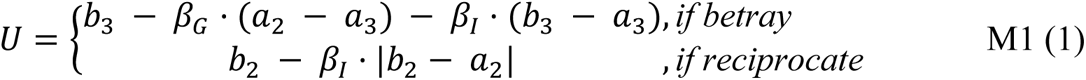

Departing from previous studies [8,17], participants were not informed of their partner’s beliefs regarding reciprocity, thus simulating real-world social interactions where individuals often lack explicit knowledge of others’ internal beliefs. The term (*a*_2_ − *a*_3_) approximates the anticipated guilt experienced upon choosing betrayal (where the partner receives *a*_3_), considering the partner’s assumed trust and expectation of reciprocity (anticipating *a*_2_), with *β*_*G*_ (0< *β*_*G*_ <10) capturing the participant’s aversion to guilt. The terms (*b*_3_ − *a*_3_) and |*b*_2_ − *a*_2_| quantify the aversion to inequity associated with betrayal and reciprocity, with *β*_*I*_ (0< *β*_*I*_ <10) representing the participant’s aversion to inequity. By design of the binary trust game, what the participant receives (*b*_3_) is constantly larger than what the partner receives (*a*_3_) in the option of betrayal, whereas the participant receives (*b*_2_) can be either greater or less than what the partner receives (*a*_2_) in the option of reciprocity. In the baseline model, inequity was quantified as the absolute difference between b_2_and a_2_ in the option of reciprocity, assuming that participants equally weigh both advantageous and disadvantageous inequity aversions [8,17]. Finally, the utility (*U*) of choosing to reciprocate and betray was entered into a SoftMax function with an inverse temperature parameter *λ* (0< *λ* <10), such that the probability of choosing to reciprocate in each trial was expressed as (**Eq.2**):

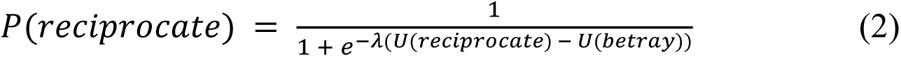

The initial model, **M1**, expanded upon the established baseline model by incorporating varying sensitivities to guilt and inequity aversion across the four experimental conditions, namely low/high trait anxiety and gain/loss framing. **M2** (**Eq. 3**) employed the Fehr–Schmidt inequity aversion model [81], which separated inequity aversion into superiority aversion (advantageous inequity aversion) and disadvantageous inequity aversion.

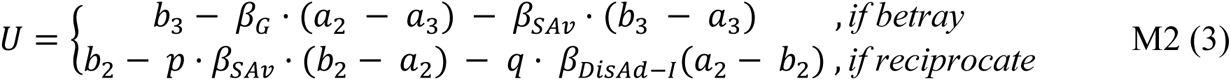

Here, *β*_*SAv*_ and *β*_*DisAd*−*I*_ represents the participant’s subjective aversion to superiority and disadvantageous inequity. *p* and *q* are conditional indicators: if *b*_2_ > *a*_2_, *p* = 1, *q* = 0; if *b*_2_ > *a*_2_, *p* = 0, *q* = 1. Given that **M2** outperformed **M1**, **M3** was developed by incorporating a reward parameter (**Eq. 4**) into **M2**, resulting in further improved model performance.

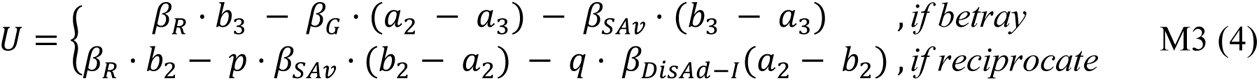

Building upon **M3**, **M4** assumes that participants exhibit a dislike for superiority in betrayal scenarios yet appreciate it in reciprocity scenarios (**Eq. 5**), leading to the replacement of the inequity aversion term in the reciprocity option with a superiority attraction component.

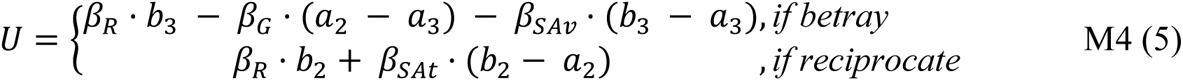

Here, the term (*b*_2_ − *a*_2_) represents the magnitude and direction of the perceived superiority (positive or negative), while and *β*_*SAt*_ reflected the participant’s sensitivity to this superiority attraction. The model comparison revealed that **M4** outperformed **M3**. Building upon **M4**, **M5** introduces a reward weighting parameter against other components (0 < *β*_*R*_ < 1) (**Eq. 6**), while **M6** assumes individuals hold a consistent trade-off between randomness and determinism in decision-making, employing a shared inverse temperature parameter (*λ*) across frames, and **M7** replaces *λ* with constant 1 as previous suggested [9]. Ultimately, model comparison confirmed **M4** as the winning model.

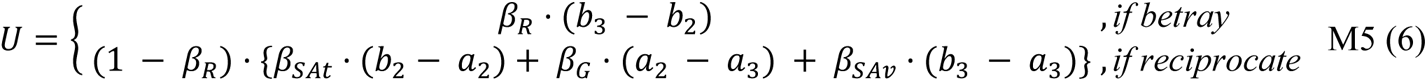

**M4** was validated using posterior predictive checks and parameter recovery [82]. For each participant and frame, model parameters were drawn and averaged from the posterior distribution to simulate decision choices and compute a predicted reciprocity rate. The correlation between these predicted and observed reciprocity rates was then used to assess the model’s predictive accuracy. Parameter recovery was conducted by refitting **M4** to the simulated dataset and assessing the correlation between the original and recovered parameters, thus validating the model’s fitting precision.

### 4.7. Statistical Analysis

All statistical analyses were performed in R (v4.1.1; www.r-project.org). LMM was performed with the lme4 package [83]. Results were considered statistically significant at the statistical threshold level *p* < .05 (two-tailed). LMM analyses were used to examine the effects of trait anxiety and context (frame) on reciprocity rate, guilt aversion, superiority aversion, reward sensitivity, and superiority attraction, as derived from the winning model. Fixed effects for trait anxiety, frame, and their interaction were included, along with subject-specific random intercepts. Differences in IRI subscales and Machiavellianism between high and low trait anxiety groups were assessed using the Mann-Whitney U test, chosen for its robustness to potential outliers and non-normal distributions often observed in these psychological measures.

To assess the contribution of each winning model component to reciprocity rate variance, a linear model was employed. Component weights were calculated using the “relaimpo” library in R. To validate the model, LMMs were used to examine the relationship between individual sensitivity to each model component and corresponding eye movement transitions. The LMM included fixed effects for individual component sensitivity, along with subject-specific random intercepts. To control for potential confounding factors, the objective value of the component, frame (gain/loss), and reaction time were included as fixed effects.

To investigate the effects of trait anxiety and frame on ERP P2, N2, and LPP amplitudes, LMMs with trial-level data were employed. The models included fixed effects for trait anxiety, frame, and their interaction, as well as subject-specific random intercepts and slopes for frame. To examine the mediating roles of P2 and LPP amplitudes, serial mediation analysis was performed using the R library “bruceR” [84], an adaptation of SPSS’s “PROCESS” [85]. Trial-level P2 and LPP amplitudes were used, along with trial-level guilt aversion and superiority aversion, calculated as the subjective value (*β*_*G*_, *β*_*SAv*_) multiplied by the corresponding objective value (a2-a3 for guilt aversion, b2-a2 for superiority aversion) [61].

To investigate the impacts of trait anxiety on the contextual effect of superiority aversion, reward sensitivity, and N2 amplitude, Mann-Whitney U tests were conducted. To assess the association between the contextual effect of superiority aversion and N2 amplitude, Spearman correlation analyses were implemented.

## 5. Additional information

## Author Contributions

**Conceptualization**: Huihua Fang.

**Data curation**: Rong Wang.

**Data arrangement:** Rong Wang.

**Formal analysis**: Huihua Fang.

**Funding acquisition**: Yuejia Luo.

**Investigation**: Huihua Fang.

**Methodology**: Huihua Fang, Zhihao Wang.

**Project administration**: Huihua Fang, Rong Wang, Qian Liu.

**Supervision**: Frank Krueger, Pengfei Xu, Yuejia Luo.

**Validation**: Huihua Fang, Frank Krueger.

**Visualization**: Huihua Fang.

**Writing – original draft**: Huihua Fang, Frank Krueger.

**Writing – review & editing**: All authors contributed to the final version.

## Data Availability Statement

The dataset and code for this manuscript are accessible on the Open Science Framework (OSF) at (<u>link</u>)

## Supplementary

### Reaction time

**Figure S1.**
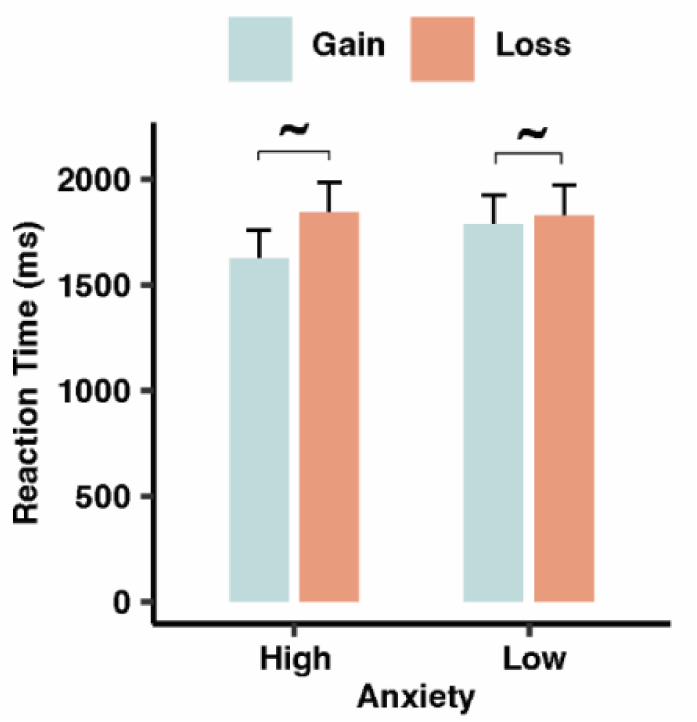
The impact of trait anxiety and context on reaction time in reciprocity decision (Mean ± SE). Individuals under Loss Frame exhibited a trend of longer reaction time than those under Gain Frame. **∼**: *p* < 0.1.

### Model prediction

**Figure S2.**
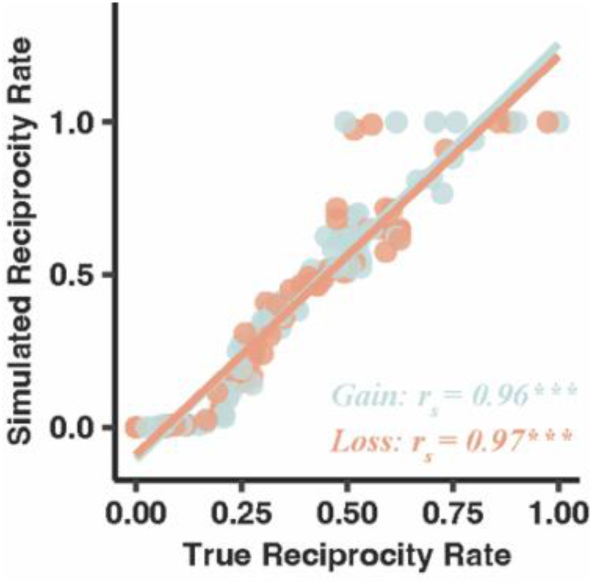
Model prediction from winning model M4. **(A)** True reciprocity rate was correlated with simulated reciprocity rate in both Gain and Loss frame. **\*\*\***: *p* < 0.001.

### Parameter recovery

**Figure S3.**
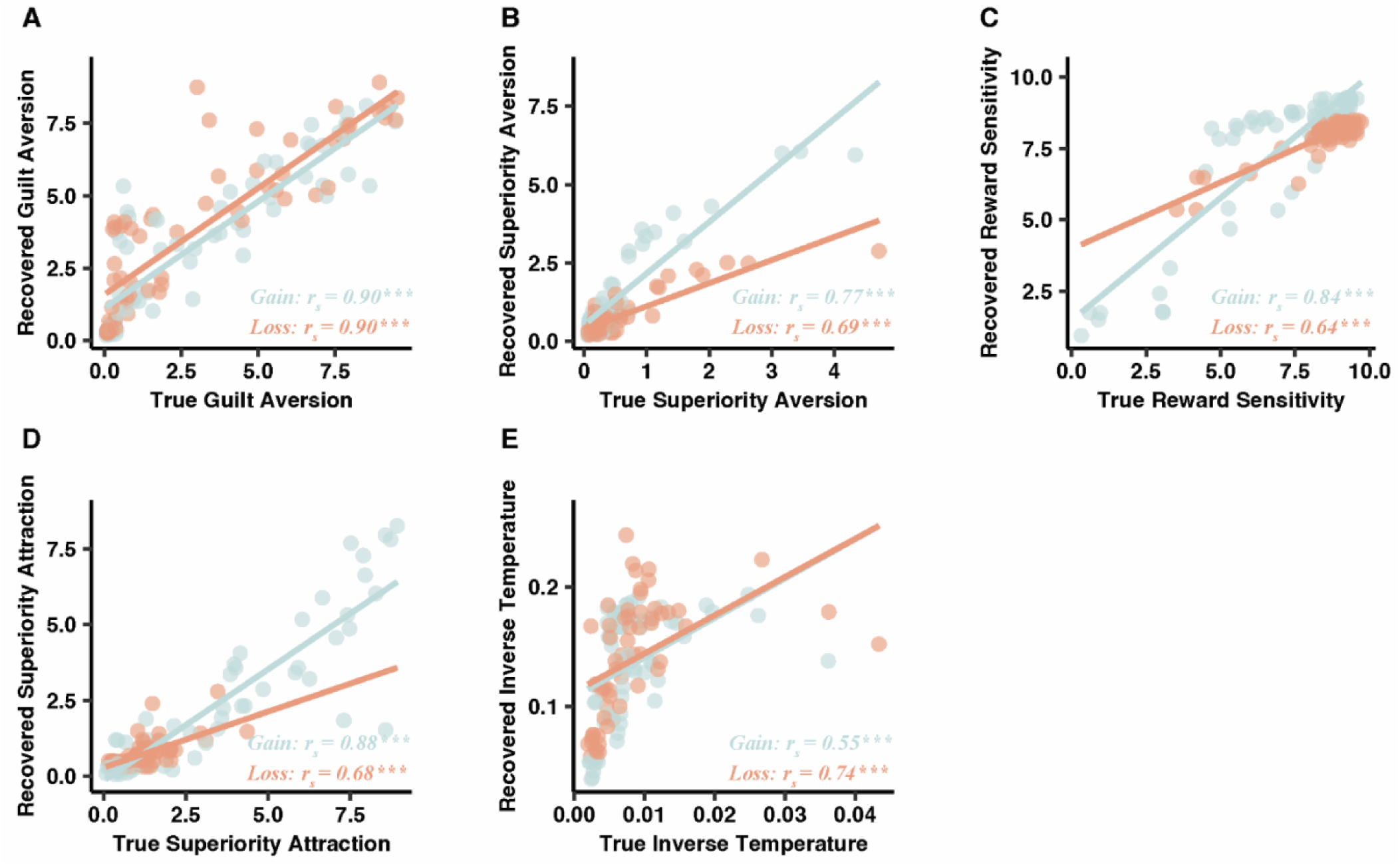
Parameter recovery from winning model M4. True parameters of **(A)** guilt aversion, **(B)** Superirority aversion, **(C)** reward sensitivity, **(D)** superirority attraction, and **(E)** inverse temperature were correlated with recovered parameters in both Gain and Loss frame. **\*\*\***: *p* < 0.001.

### Displaying version effect

The LMMs revealed no significant main effect of displaying version on the recirpocity rate (χ²(1) = 1.64, *p* = 0.650) and parameters of guilt aversion (χ²(1) = 0.52, *p* = 0.915), superiority aversion (χ²(1) = 2.62, *p* = 0.453), reward sensitivity (χ²(1) = 2.73, *p* = 0.435), superiority attraction (χ²(1) = 3.16, *p* = 0.368) and inverse temperature (χ²(1) = 0.758, *p* = 0.860) from the winning model **M4**.

